# Dominance of ST131 *Escherichia coli* carrying *bla*_CTX-M_ in patients with bloodstream infections caused by cephalosporin-resistant strains in Australia, New Zealand and Singapore: whole genome analysis of isolates from a randomised trial

**DOI:** 10.1101/181768

**Authors:** Patrick N. A. Harris, Nouri L. Ben Zakour, Leah W. Roberts, Alexander M. Wailan, Hosam M. Zowawi, Paul A. Tambyah, David C. Lye, Roland Jureen, Tau H. Lee, Mo Yin, Ezlyn Izharuddin, David Looke, Naomi Runnegar, Benjamin Rogers, Hasan Bhally, Amy Crowe, Mark A. Schembri, Scott Beatson, David L. Paterson, on behalf of the MERINO Trial investigators

**Author notes:** Corresponding author: Dr. Patrick N. A. Harris, University of Queensland Centre for Clinical Research, Building 71/918 Royal Brisbane & Women’s Hospital Campus, Herston, QLD, 4029.

## Abstract

**Objectives:** To characterise multi-drug resistant *Escherichia coli* isolated from patients in Australia, New Zealand and Singapore with bloodstream infection (BSI).

**Methods:** We prospectively collected third-generation cephalosporin resistant (3GC-R) *E. coli* from blood cultures obtained from patients enrolled in a randomised controlled trial. Whole genome sequencing was used to characterise antibiotic resistance genes, sequence types (STs), plasmids and phylogenetic relationships. Antibiotic susceptibility was determined using disk diffusion and Etest.

**Results:** A total of 70 *E. coli* were included, of which the majority were ST131 (61.4%). BSI was most frequently from a urinary source (69.6%), community-associated (62.9%) and in older patients (median age 71 years [IQR 64-81]). The median Pitt bacteraemia score at presentation was 1 (IQR 0-2, range 0-3) and ICU admission was infrequent (3.1%). ST131 possessed significantly more acquired resistance genes than non-ST131 (p=0.003). Clade C1/C2 ST131 predominated (30.2% and 53.5% of all ST131 respectively) and these were all resistant to ciprofloxacin. All clade A ST131 were community-associated. The predominant ESBL types were *bla*_CTX-M_ (78.6% of isolates) and were strongly associated with ST131, with the majority *bla*_CTX-M-15_. Clade C1 was associated with *bla*_CTX-M-14_ and *bla*_CTX-M-27_, whereas *bla*_CTX-M-15_ predominated in clade C2. Plasmid-mediated AmpC (p-AmpC) genes (mainly *bla*_CMY-2_) were also frequent (17.1%) but were more common with non-ST131 strains (p< 0.001). The majority of plasmid replicon types were IncF.

**Conclusions:** In a prospective collection of 3GC-R *E. coli* causing BSI in the Australasian region, community-associated Clade C1/C2 ST131 predominate in association with *bla*_CTX-M_ ESBLs, although a significant proportion of non-ST131 strains carried *bla*_CMY-2_.

## Introduction

In recent decades, resistance to beta-lactam antibiotics in Enterobacteriaceae has become increasingly common. Of particular concern has been the rising prevalence of resistance to 3^rd^ generation cephalosporins (3GCs), which includes key agents such as ceftriaxone, cefotaxime or ceftazidime.^1-3^ This phenomenon has largely arisen from the dissemination of genes encoding extended-spectrum beta-lactamase (ESBL) or, less frequently, plasmid-mediated AmpC (p-AmpC) enzymes.^4-6^ These resistance genes are often acquired by plasmid transfer and may be associated with other antibiotic resistance determinants, rendering organisms multi-drug resistant (MDR).^7^ The global emergence of infections caused by ESBL-producing *E. coli*, in both the community and hospital setting, has been driven by the acquisition of CTX-M-type ESBL genes (especially *bla*_CTX-M-15_) in the successful pandemic clone of *E. coli*, sequence type 131 (ST131).^8-11^ *E. coli* ST131 belong to the B2 phylogenetic subgroup I, and are mostly serotype O25b:H4.^12^ Within ST131, further sublineages have been delineated according to *fimH* (type 1 fimbrial adhesin, FimH) alleles, phylogenetic clades (A, B, C1 and C2) and associated resistance genes.^8, 13^ A globally dominant fluoroquinolone-resistant *fimH*30 sub-clonal lineage, defined as *H*30-R (to differentiate from the ancestral fluoroquinolone susceptible *H*30 strains) or clade C, has been described.^8, 14^ *H*30-R/clade C strains contain fluoroquinolone resistance mutations in the chromosomal *gyrA* and *parC* genes^15^ and have been associated with poor clinical outcomes.^16^ Within this sub-lineage, a pathogenic ST131 subclone containing *bla*_CTX-M-15_ has been referred to as *H*30-Rx^14^ or clade C2.^8^ Specific Incompatibility (Inc) F-type plasmids have also been described in association with fluoroquinolone-resistant ST131-*H*30 clades, with IncF type F1:A2:B20 plasmids associated with the *H*30-R/C1 clade and IncF type F2:A1:B-plasmids associated with the *H*30-Rx/C2 clade.^17-19^

Resistance to beta-lactams mediated by ESBLs drives the use of broader spectrum antibiotics such as carbapenems (e.g. meropenem)^4^, providing selection pressure for carbapenem-resistance in gram-negative bacteria. Of great concern has been the emergence of transmissible carbapenemases in common Enterobacteriaceae.^20^ As part of an international randomised controlled trial of piperacillin-tazobactam (a potential ‘carbapenem-sparing’ agent) versus meropenem for the treatment of 3GC-resistant *E. coli* causing bloodstream infection,^21^ we aimed to further characterise these isolates using whole genome sequencing.

### Objectives

To analyse a prospectively collected series of ceftriaxone non-susceptible *E. coli* isolated from blood cultures in patients in Australia, New Zealand and Singapore using whole genome sequencing, in order to characterise antibiotic resistance genes, sequence types, phylogenetic relationships and plasmid structures.

## Methods

### Bacterial isolates and clinical data

*E. coli* isolates were collected prospectively during an international multi-centre randomised trial comparing treatment options for bloodstream infections caused by ceftriaxone-resistant *E. coli* or *Klebsiella* spp. (the ‘MERINO’ trial: Australian New Zealand Clinical Trials Register (ANZCTR), Ref no: ACTRN12613000532707 and the U.S. National Institute of Health ClinicalTrials.gov register, Ref no: NCT02176122).^21^ All *E. coli* blood culture isolates were included from 70 patients enrolled in the trial for the first 18 months of recruitment (from February 2014 until August 2015) from 8 hospital sites in 3 countries. To be eligible for the trial, patients had to have at least one monomicrobial blood culture growing *E. coli*, with resistance to ceftriaxone, ceftazidime or cefotaxime determined by methods used in the local laboratories (all of which were internationally accredited). All blood culture isolates were stored at the recruiting site laboratory at −80°C in cryovials containing glycerol and nutrient broth and later shipped to the co-ordinating laboratory in Queensland, Australia. For the purposes of this study, only the first *E. coli* isolated from blood cultures for each enrolled patient were included in the genomic analysis. Relevant clinical data were collected and managed using the REDCap^22^ electronic data capture tool hosted at the University of Queensland. Ethics approval for the study was provided by the Royal Brisbane and Women’s Hospital (Ref: HREC/12/QRBW/440), the National Healthcare Group (NHG) Domain Specific Review Board (DSRB) in Singapore (NHG DSRB Ref: 2013/00453) and the New Zealand Health and Disability Ethics Committee (Ref: 14/NTB/52).

### Phenotypic susceptibility testing

All isolates were tested at the co-ordinating laboratory against a standard panel of antibiotics used to treat gram-negative infections by disk diffusion according to EUCAST standards.^23^ Agents tested included ceftriaxone, ceftazidime, cefepime, cefoxitin, aztreonam, ertapenem, gentamicin, amikacin, ciprofloxacin, co-trimoxazole, and amoxicillin-clavulanate. In addition, minimum inhibitory concentrations (MICs) for piperacillin-tazobactam, meropenem (the two comparator drugs used in the trial) and ceftriaxone were determined by Etest (bioMérieux). ESBL production was confirmed using combination disk testing with ceftriaxone and ceftazidime with and without clavulanate; an increase in zone diameter ≥5mm with the addition of clavulanate confirmed ESBL production.^24^

### DNA extraction and library preparation

After sub-culture onto LB agar to check for viability and purity, genomic DNA was extracted using the MoBio Ultrapure kit and quantified by spectrophotometry (NanoDrop; ThermoFisher) and fluorometry (Qubit; ThermoFisher). Paired-end DNA libraries were prepared using the Illumina Nextera kit in accordance with the manufacturer’s instructions.

### Whole genome sequencing

Whole genome sequencing was performed in two batches using Illumina HiSeq (100 bp paired end) and MiSeq (300 bp paired end) at the Australian Genome Research Facility (AGRF), University of Queensland, St Lucia. MiSeq raw reads were trimmed conservatively to 150 bp and filtered using Nesoni (v0.130) to remove Illumina adaptor sequences, reads shorter than 80 bp and bases below Phred quality 5 (https://github.com/Victorian-Bioinformatics-Consortium/nesoni). Strains were checked for contamination using Kraken (0.10.5-beta) as implemented through Nullarbor (default settings).^25^

### Resistance gene detection, MLST and Plasmid typing

Antibiotic resistance genes were detected by using Abricate (v0.2) with the ResFinder database against SPAdes assemblies (v3.6.2) as implemented through the pipeline analysis tool Nullarbor (default settings).^25^ Multi-locus sequence typing was undertaken using the mlst tool as implemented through Nullarbor. Plasmid replicon typing and plasmid multilocus typing for IncF plasmids were performed using PlasmidFinder and pMLST.^26^

### Fluoroquinolone resistance SNP detection

Filtered reads were mapped to the complete ST131 *E. coli* reference strain EC958 (Genbank: HG941718.1) using Bowtie as implemented through Nesoni. Non-synonymous mutations were identified using Nesoni nway and manually compared to known mutations in *gyrA* and *parC* associated with quinolone resistance.^27, 28^

### Phylogenetic analysis

Reads for all isolates (n=70) were mapped to the complete ST131 reference EC958 (Genbank: HG941718.1)^29^ using Nesoni under default settings. Single Nucleotide Polymorphisms (SNPs) identified between isolates and the reference EC958 were used to create pseudogenomes for each isolate by substituting the relevant SNPs into the EC958 chromosome using an in-house script. Multiple sequence alignment of the pseudogenomes was used as input for Gubbins (v1.3.4)^30^ using the (GTR)GAMMA substitution model to parse recombinant regions. The remaining 211,920 SNPs were used to generate a phylogenetic tree using RAxML (8.2.9)^31^ under the (GTR)GAMMA substitution model with Lewis ascertainment bias correction and a random seed of 456 (100 bootstraps). An ST131 only tree (n=43) was also created in the same manner using 2,248 recombination-free SNPs and 1000 bootstraps. Phylogenetic trees and associated meta-data were visualised using Evolview.^32^

### Statistical tests

Data describing patient demographics, phenotypic susceptibility, clinical variables and genotypic data for all cases were tabulated, with proportions expressed as percentages and median, mean or inter-quartile ranges calculated as appropriate for scale variables. Categorical variables were compared using Pearson’s χ^2^ test. Comparisons of mean values in normally distributed data were compared using the t-test. The Mann-Whitney U test was used for non-parametric data. Statistical analysis was performed using Stata v.13.1 (StataCorp; TX, USA) and graphical images prepared using Prism v.7 (GraphPad Software; CA, USA). A p-value <0.05 was considered significant.

## Results

A total of 70 *E. coli* bloodstream isolates were included. The background clinical and demographic details of enrolled patients are summarised in Table 1. The source of bloodstream infection was most frequently the urinary tract (48/70, 69.6%) and infections were mostly community-associated (44/70, 62.9%). There was a predominance of patients reporting Chinese ethnicity (38/70, 54.3% of all cases), reflecting the demographics of the largest recruiting sites in Singapore. There were also a greater proportion of male patients (60%), but this was not statistically significant (p=0.12). Patients tended to be older (median age 71, IQR 64-81 years, range 20 to 94 years), although only a small proportion (5.8%) were admitted from nursing homes. The majority of patients had less severe acute illness (median Pitt score 1, IQR 0 to 2, range 0-3; where a score ≥4 reflects the presence of critical illness with high mortality^33^) and relatively low co-morbidity scores (Charlson score median 2, IQR 1 to 4, range 0 to 11) and were infrequently admitted to the ICU (3.1%).

**Table 1:** Baseline clinical and demographic variables

| Factor | Level | N (%) |
| --- | --- | --- |
| **Region** | Singapore | 44 (63%) |
|  | Brisbane (AUS) | 13 (19%) |
|  | Melbourne (AUS) | 8 (11%) |
|  | Auckland (NZ) | 5 (7%) |
| **Age [years], mean (SD)** |  | 69.9 (15.2) |
| **Gender** | Female | 28 (40%) |
|  | Male | 42 (60%) |
| **Source of Bacteraemia** | Urinary tract infection | 48 (70%) |
|  | Intra-abdominal infection | 8 (12%) |
|  | Pneumonia | 1 (1%) |
|  | Other | 9 (13%) |
|  | Unknown | 3 (4%) |
| **Acquisition** | Community-associated | 44 (63%) |
|  | Healthcare-associated | 26 (37%) |
| **Pitt score, median (IQR)** |  | 1 (0, 2) |
| **Charlson Score, median (IQR)** |  | 2 (1, 4) |
| **Any CTX-M ESBL** |  | 55 (79%) |
| **AmpC β-lactamase** |  | 12 (17%) |
| **Surgery within 14 days** |  | 5 (7%) |
| **Central venous catheter** |  | 6 (9%) |
| **Immune suppression** |  | 10 (14%) |
| **ICU admission** |  | 2 (3%) |
| **Nursing home resident** |  | 4 (6%) |
| **Total** |  | **70** |

IQR = inter-quartile range, SD = standard deviation, AUS = Australia, NZ = New Zealand, ICU = intensive care unit

Strains demonstrated a variable antibiogram according to ST131 clade (see Table 2), but were frequently resistant to trimethoprim-sulphamethoxazole (46/70, 65.7%) or fluoroquinolones (52/70, 74.3%). There was universal resistance to ciprofloxacin in clade C1/C2 ST131, compared with only 50% and 48.2% in clade A and non-ST131 strains respectively (p<0.001). Resistance to aminoglycosides was variable, with (25/70, 35.7%) testing resistant to gentamicin, but none were resistant to amikacin. By MIC testing, 97.1% (68/70) were susceptible to piperacillin-tazobactam (median 2mg/L, range 1-24mg/L, IQR 1.5-4; EUCAST breakpoint for susceptibility ≤8 mg/L^34^) (see supplementary figure 1). All strains were susceptible to meropenem (MIC_90_ = 0.047 mg/L; median 0.023 mg/L, range 0.012-0.19 mg/L; EUCAST breakpoint for susceptibility ≤2 mg/L^34^), although one strain (MER-86) was non-susceptible to ertapenem. The majority (90.1%) demonstrated ceftriaxone MICs ≥32 mg/L (range 0.064 to ≥32 mg/L; median ≥32 mg/L, MIC_90_ and MIC_50_ ≥32 mg/L). Two strains, which were susceptible to ceftriaxone by MIC, were resistant to ceftazidime. Phenotypic resistance to third-generation cephalosporins could not be detected in one strain (MER-34) when retested in the co-ordinating laboratory, although it was found to posses TEM-176 ESBL by sequencing.

**Table 2:** Antibiotic resistance profile of *E. coli* strains according to ST-131 clade

| Clade | N | Cephalosporin |  |  |  | BLBLI |  | Carbapenem |  | Monobactam | Sulphonamide | Aminoglycoside |  | Quinolone |
| --- | --- | --- | --- | --- | --- | --- | --- | --- | --- | --- | --- | --- | --- | --- |
|  |  | CTX | CAZ | FEP | FOX | AMC | PTZ | MEM | ETP | ATM | SXT | GM | AK | CIP |
|  |  | Non-susceptible N (%) |  |  |  |  |  |  |  |  |  |  |  |  |
| A | 6 | 5<br>(83) | 5<br>(83) | 5<br>(83) | 1<br>(17) | 4<br>(67) | 0<br>(0) | 0<br>(0) | 0<br>(0) | 5<br>(83) | 4<br>(67) | 3<br>(50) | 0<br>(0) | 3<br>(50) |
| B | 1 | 1<br>(100) | 1<br>(100) | 1<br>(100) | 0<br>(0) | 1<br>(100) | 0<br>(0) | 0<br>(0) | 0<br>(0) | 1<br>(100) | 1<br>(100) | 1<br>(100) | 0<br>(0) | 0<br>(0) |
| C1 | 13 | 13<br>(100) | 4<br>(31) | 12<br>(92) | 1<br>(8) | 5<br>(38) | 0<br>(0) | 0<br>(0) | 0<br>(0) | 12<br>(92) | 8<br>(62) | 4<br>(31) | 0<br>(0) | 13<br>(100) |
| C2 | 23 | 23<br>(100) | 21<br>(91) | 22<br>(96) | 1<br>(4) | 21<br>(91) | 2<br>(9) | 0<br>(0) | 0<br>(0) | 23<br>(100) | 19<br>(83) | 13<br>(57) | 0<br>(0) | 23<br>(100) |
| Non-ST131 | 27 | 25<br>(93) | 22<br>(81) | 14<br>(52) | 10<br>(37) | 20<br>(74) | 0<br>(0) | 0<br>(0) | 1<br>(4) | 21<br>(78) | 14<br>(52) | 4<br>(15) | 0<br>(0) | 13<br>(48) |
| All | 70 | 67<br>(96) | 53<br>(76) | 54<br>(77) | 13<br>(19) | 51<br>(73) | 2<br>(3) | 0<br>(0) | 1<br>(1) | 62<br>(89) | 46<br>(66) | 25<br>(36) | 0<br>(0) | 52<br>(74) |

BLBLI = beta-lactam/beta-lactamase inhibitor, CTX=ceftriaxone, CAZ=ceftazidime, FEP=cefepime, FOX=cefoxitin, AMC=amoxicillin-clavulanate, PTZ=piperacillin-tazobactam, MEM=meropenem, ETP=ertapenem, ATM=aztreonam, SXT=trimethoprim-sulphamethoxazole, GM=gentamicin, AK=amikacin, CIP=ciprofloxacin.

**Figure 1:**
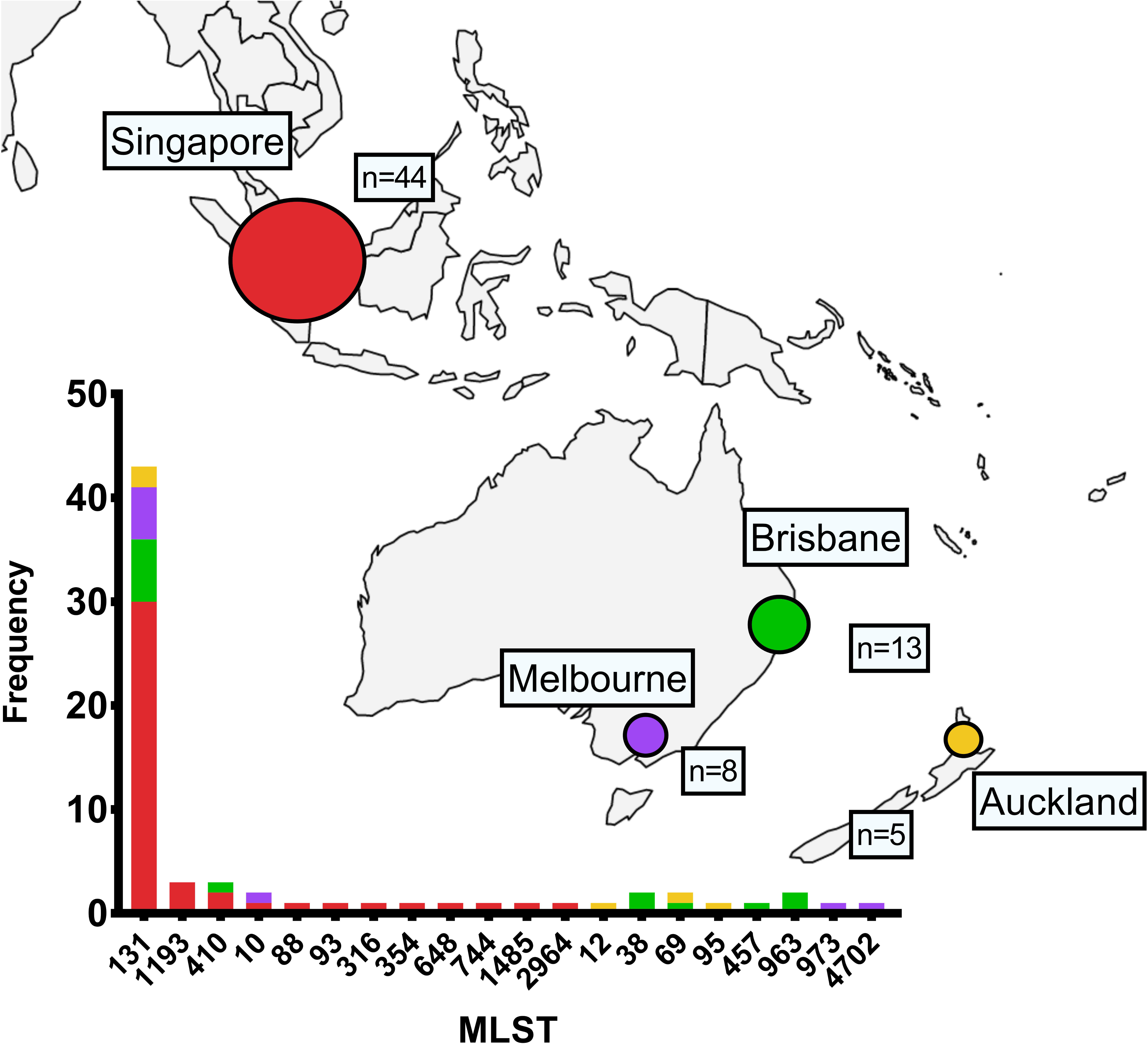
*In silico* MLST of ESBL or AmpC-producing *E. coli* isolated from blood, by region

### MLST and phylogenetic grouping of ST131

A clear predominance of strains were ST131 by *in silico* MLST (43/70, 61.4%), with other strains broadly distributed across a number of other STs (figure 1). The majority of ST131 strains belonged to clades C1 (30.2%) or C2 (53.5%), with strains from clades B (2.3%) and A (14.0%) seen less frequently.

### SNPs and phylogenetic relationships of ST131

All clade A ST131 were community-associated, with a mixture of community and healthcare-associated infections observed for strains in clades C1 and C2. There was evidence of clustering of closely related strains within certain hospitals (e.g. MER-27/25 in Hospital E; MER-8/10 and MER-37/39 in Hospital A; MER-78/79 in Hospital B; MER-65/66 in Hospital G) (figure 2), although these represented both community and healthcare-associated infections. It is also notable that closely related strains were identified in different countries, emphasising the global dissemination of ST131. A phylogenetic tree of all *E. coli* strains can be found in supplementary figure 2.

**Figure 2:**
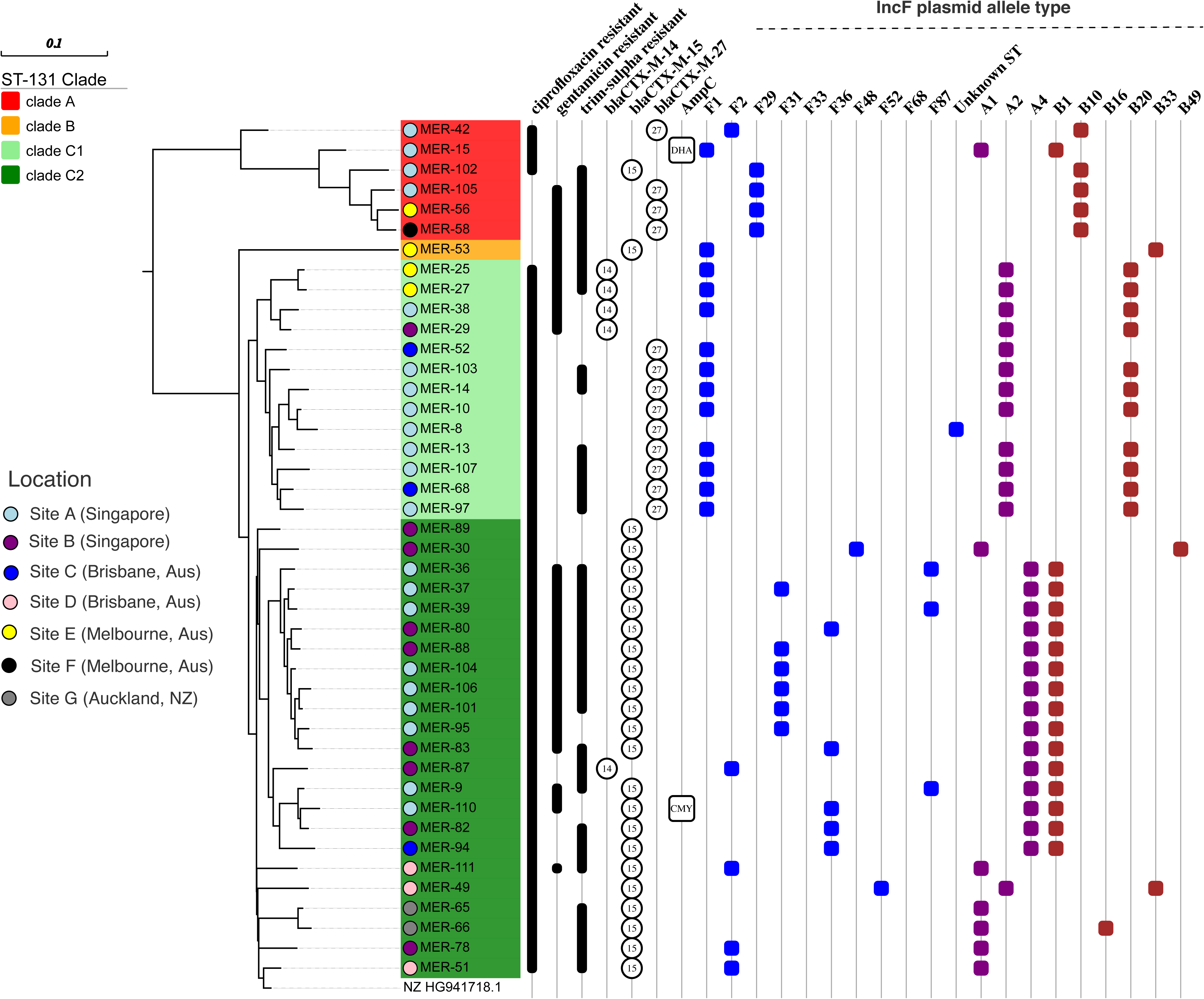
Phylogenetic tree of ST131 *E. coli* based on core genome SNPs; clade, antibiotic resistance, ESBL/p-AmpC type and IncF plasmid type are shown Aus = Australia; NZ = New Zealand

### Resistance genes

The median number of acquired resistance genes detected for each isolate was 9. One strain (MER-90) possessed a total of 17 acquired resistance genes, including beta-lactamases (*bla*_CMY-2_, *bla*_TEM-1B_), aminoglycoside resistance genes (*aac(3)-IId*-like, *aadA1*-like, *aadA2*, *aph(3’)-Ic*-like, *strA*, *strB*), resistance genes related to folate metabolism (*dfrA12*, *dfrA14-like*), fluoroquinolones (*qnrS1*), sulphonamides (*sul1, sul2, sul3*), tetracyclines (*tet(A)*), phenicols (*floR-like*) and macrolides (*mph(A)*). The number of acquired antibiotic resistance genes was significantly greater in ST131 than non-ST131 strains (p=0.003) (figure 3A). However, the number of resistance genes did not vary significantly across ST131 clades (supplementary figure 3). The complete distribution of resistance genes can be found in supplementary dataset 1.

**Figure 3:**
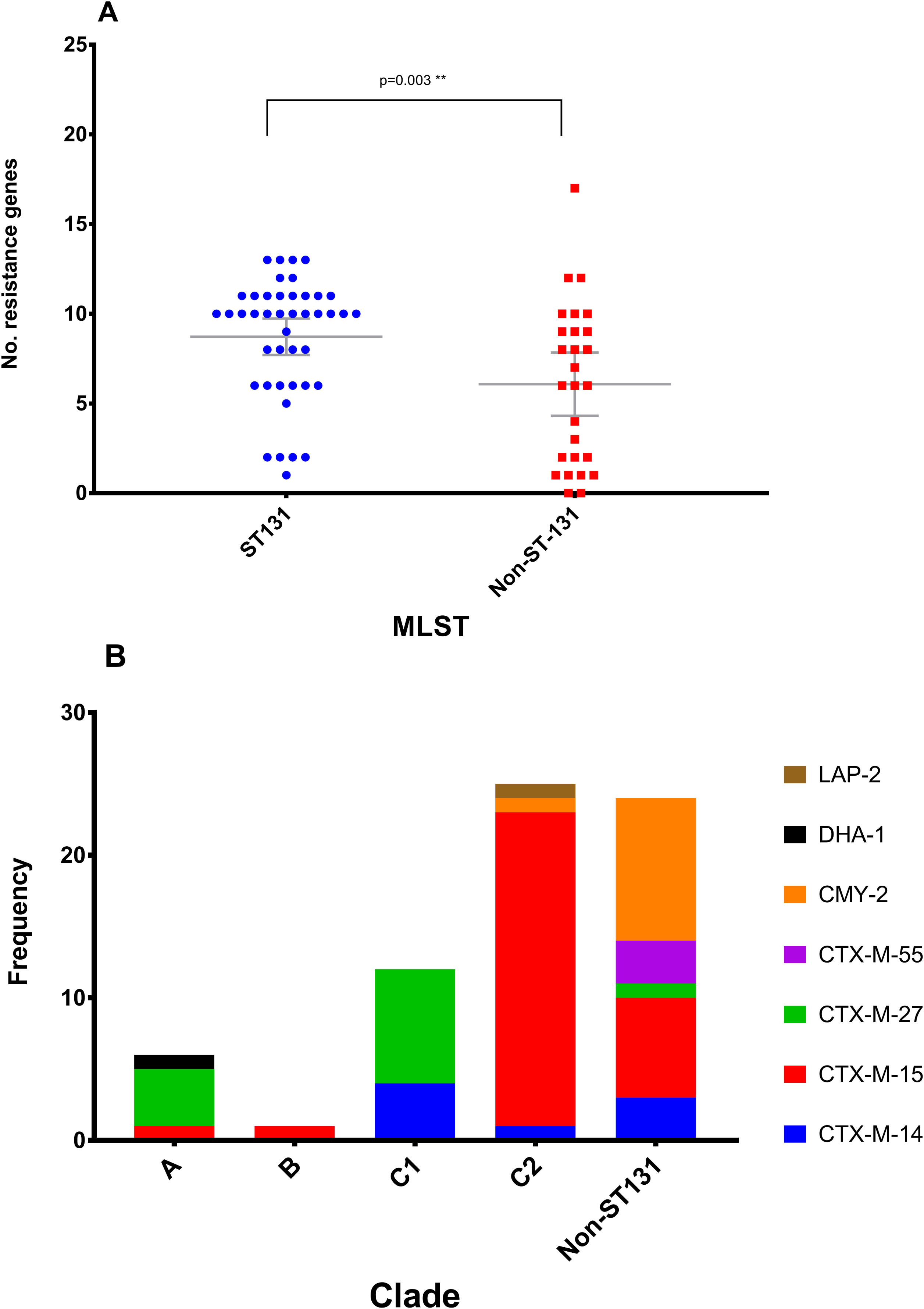
Panel A | Number of resistance genes by sequence type; **Panel B** | Distribution of ESBL and p-AmpC genes across ST131 clades (A, B, C1 and C2) and non-ST131 *E. col* Only acquired resistance genes detected by whole genome sequencing are shown. MLST = *in silico* multi-locus sequence type, grey bars show means with 95% confidence intervals. Groups were compared using Mann-Whitney U-test; ** significant at p< 0.005 level

### Beta-lactamases

The predominant ESBL genes identified were *bla*_CTX-M_, seen in 78.6% (55/70) of isolates. The presence of *bla*_CTX-M_ was strongly associated with ST131; 95% of ST131 possessed *bla*_CTX-M_ ESBLs, compared with only 56% of non-ST131 (p< 0.001) (Table 3). These were either from CTX-M group 9 (*bla*_CTX-M-14,_ *bla*_CTX-M-27_) or CTX-M group 1 (*bla*_CTX-M-15,_ *bla*_CTX-M-55_)^35^, with the majority *bla*_CTX-M-15_. Clade C1 ST131 were associated with *bla*_CTX-M-14_ and *bla*_CTX-M-27_, whereas *bla*_CTX-M-15_ predominated in clade C2 (figure 3B). Two strains from Singapore (MER-33, MER-34) possessed a TEM-variant ESBL (*bla*_TEM-176_), of which one was co-harboured with *bla*_CMY-2_ (MER-33). No SHV-group ESBLs were identified in these *E. coli* isolates.

**Table 3:** Presence of acquired resistance genes by sequence type

| Resistance genes | All strains | ST131 | Non-ST131 | p-value |
| --- | --- | --- | --- | --- |
| **CTX-M-type ESBL** | 56 (79%) | 41 (95%) | 14 (52%) | **<0.001** |
| *CTX-M-14* | 8 (11%) | 5 (12%) | 3 (11%) | 0.95 |
| *CTX-M-15* | 31 (44%) | 24 (56%) | 7 (26%) | **0.014** |
| *CTX-M-27* | 14 (20%) | 13 (30%) | 1 (4%) | **0.007** |
| *CTX-M-55* | 3 (4%) | 0 (0%) | 3 (11%) | **0.025** |
| **Acquired AmpC β-lactamase** | 12 (17%) | 2 (5%) | 10 (37%) | **<0.001** |
| **Aminoglycoside modifying enzymes** | 53 (76%) | 37 (86%) | 16 (59%) | **0.011** |
| **Acquired quinolone resistance** | 8 (11%) | 2 (5%) | 6 (22%) | **0.025** |
| **Folate pathway resistance** | 38 (54%) | 32 (74%) | 6 (22%) | **<0.001** |
| **Sulphonamide resistance** | 48 (69%) | 35 (81%) | 13 (48%) | **0.004** |
| **Tetracycline resistance** | 39 (56%) | 23 (53%) | 16 (59%) | 0.64 |
| **Total** | **70** | **43** | **27** |  |

The second most common group of beta-lactamases with the ability to hydrolyse third-generation cephalosporins were acquired AmpC beta-lactamase genes, found in 17.1% (12/70). The presence of acquired AmpC was not clearly associated with specific STs, but was more common in non-ST131 strains (37.0% vs 4.7%; p< 0.001). These were predominantly *bla*_CMY-2_, although a single strain carried *bla*_DHA-_ 1. A single clade C2 ST131 strain from Singapore possessed *bla*_CMY-2_ (in association with *bla*_CTX-M-15_) (figure 2). Although *bla*_CMY-2_ are usually acquired on plasmids, in 4 strains (3 from Australia [MER-2, MER-4, MER-43] and 1 from Singapore [MER-99]) there was evidence to suggest chromosomal integration. However, due to repetitive regions surrounding the *bla*_CMY-2_ region, the complete context could not be ascertained in all isolates using short read sequencing alone. Two of the Australian CMY-2-producing strains [MER-2 and MER-4] were of the same ST and near identical on SNP analysis suggesting a common exposure source or direct transmission within the healthcare setting, given that their admissions overlapped in time (but on separate wards). Further details of the genetic context of *bla*_CMY-2_ and *bla*_DHA-1_ can be found in supplementary figures 4, 5, 6 and 7.

Other narrow-spectrum beta-lactamases such as *bla*_OXA-1_ or *bla*_TEM-1B_ were common (seen in 30.0% and 34.3% respectively). A single strain [MER-89] possessed *bla*_LAP-2_, and two [MER-86 and MER-110] carried both *bla*_CTX-M_ and *bla*_CMY_. No carbapenemase genes were identified. In one strain [MER-100], resistance to ceftazidime (but not ceftriaxone) was not clearly explained by resistance gene profiling, with no beta-lactamase genes identified, although an altered −35 box (TTGACA) was found in its promotor, which has been associated with overexpression of the *ampC* promotor.^36^ A single strain [MER-86] demonstrated resistance to ertapenem and was found to have disruption in *ompF*, which has been associated with reduced susceptibility to ertapenem when associated with broad spectrum beta-lactamases.^37^

### Aminoglycoside resistance genes

The presence of aminoglycoside modifying enzymes (AMEs) was common (seen in 76%) and was encountered more frequently in ST131 (86% vs 59%; p=0.011). There were a variety of AME types identified, including those belonging to the *aadA*, *aac(3)*, *aph(3’)* groups, as well as streptomycin resistance genes *strA* and *strB.* No genes encoding 16S methylase enzymes (e.g. *arm*, *rmt*), which mediate broad class resistance to aminoglycosides, were detected.

### Fluoroquinolone resistance genes

Acquired quinolone resistance genes (i.e. those not mediated by SNPs in regions associated with quinolone-resistance) were seen in 11% (8/70). These genes included *qnrS1*, *qnrB4*, *qnrB66*-like, *oqxA* or *aac(6’)Ib-cr* (which also mediates aminoglycoside resistance in addition to low-level quinolone resistance). The presence of these genes was more commonly seen in non-ST131 than ST131 (22% vs 5%; p=0.025).

All clade C strains (and a single clade A strain [MER-42]) were identical to the EC958 reference strain with respect to mutations in *parC* and *gyrA* (supplementary tables 1 and 2). Phenotypic ciprofloxacin resistance was largely congruent with the presence of SNPs in *parC* and *gyrA* known to be associated with fluoroquinolone resistance, or the presence of acquired quinolone resistance determinants. However, in a handful of strains (e.g. MER-34, MER-26) phenotypic ciprofloxacin resistance was not evident despite the presence of acquired resistance genes (e.g. *qnrS1*, *aac(6’)Ib-cr* or *oqxA/B*). This may reflect the limited sensitivity of disc diffusion methods to detect low-level quinolone resistance. Certain *gyrA* SNPs (e.g. 83L) were not by themselves associated with phenotypic ciprofloxacin resistance unless accompanied by additional SNPs (e.g. 87N or 87Y) (supplementary table 2).

### Sulphonamide and folate pathway resistance genes

Sulphonamide resistance genes (*sul1*, *sul2* or *sul3*) were common, and present in 69% (48/70) of strains, as were folate synthesis pathway (e.g. trimethoprim) resistance genes (54%, 38/70), such as *dfrA1*, *dfrA7*, *dfrA12*, *dfr14* and *dfrA17*. The presence of sulphonamide resistance and trimethoprim resistant genes were more common in ST131 (81% vs 48%, p=0.004, and 74% vs 22%, p< 0.001, respectively).

### Other resistance genes

Genes mediating resistance to tetracyclines (specifically *tet(A)* and *tet(B)*) were seen in 56% (39/70), but were equally distributed between ST131 and non-ST131. Other frequently identified resistance genes included those mediating resistance to chloramphenicol (e.g. *catA*, *florR*) or macrolides (e.g. *mph(A)*).

### Plasmids

The majority of plasmid replicon types were identified as IncF. According to the PubMLST scheme (www.pubmlst.org/plasmid), plasmids seen in clade C1 ST131 were mainly IncF plasmid type F1:A2:B20 (76.9%), with the remainder IncF type F1:A2:B- or IncI1 types [ST-79 or unknown ST]. Amongst clade C2 ST131, IncF types F31:A4:B1 or F36:A4:B1 were most common (22.7% and 27.3% respectively), with IncF type F2:A1:B-plasmids only seen in 18.2% (Figure 2). Only three clade C2 strains contained IncI1 or IncN plasmids. The full description of plasmid replicon types detected is provided in supplementary dataset 1.

## Discussion

This prospective collection of ESBL and p-AmpC-producing *E. coli* bloodstream isolates provides insight into the current clinical and molecular epidemiology of these infections within Australia, New Zealand and Singapore. The clear predominance of ST131 carrying CTX-M-type ESBLs is striking and reflects how this pandemic clone has emerged as a highly successful human pathogen. As has been described elsewhere, CTX-M-type ESBLs have now displaced TEM- or SHV-type ESBLs in many parts of the world^10, 38^, and the latter were not seen in this contemporary collection of *E. coli* bloodstream isolates. It is also notable that the majority of cases were of community-onset, with their origin from the urinary tract and in patients over the age of 65 years. This also reflects the shifting epidemiology, whereby an increasing proportion of infections caused by ESBL-producing *E. coli* are community acquired.^39^ This contrasts to previous decades, following the first description of ESBLs, where nosocomial acquisition was common and TEM and SHV-type EBSLs predominated.^4^

Different beta-lactamase genes were associated with certain *E. coli* lineages. As has been described previously, *bla*_CTX-M-15_ was largely restricted to the C2 clade amongst ST131, whereas *bla*_CTX-M-27_ and *bla*_CTX-M-14_ were found in clade C1.^8, 40^ A second notable finding is the emerging prevalence of 3GC-R *E. coli* with acquired AmpC as a cause of bloodstream infections; these were the second most commonly encountered broad-spectrum beta-lactamase after CTX-M-type ESBLs in this cohort. Having been previously under-appreciated, p-AmpC enzymes are increasingly recognised as a prominent mediator of resistance in *E. coli*.^41-45^ In this cohort, p-AmpC (mainly *bla*_CMY-2_) were not associated with ST131 or any other ST. Previous studies have demonstrated the predominant p-AmpC enzyme amongst *E. coli* has been CMY-2^41, 44, 46^, with evidence that *bla*_CMY-2_ has been mobilised from the *Citrobacter freundii* chromosome in association with IS*Ecp1.*^47^ IS*Ecp1* was identified in all but two of the *E. coli* strains carrying *bla*_CMY-2_ in our collection, with these associated with IS*1294* and a truncated IS*Ecp1* (supplementary figure 5).

IncF type plasmids have a host range that is limited to Enterobacteriaceae and contribute to bacterial fitness via antibiotic resistance and virulence determinants.^48^ These plasmids have been associated with the rapid emergence and global spread of *bla*_CTX-M-15_, as well as genes encoding resistance to aminoglycosides and fluoroquinolones (e.g. *aac(6’)-Ib-cr*, *qnr*, *armA*, *rmtB*).^48, 49^ Similar patterns were also seen in this study, where the majority of plasmids were of IncF type. There was an association between *bla*_CTX-M-15_, *bla*_OXA-1_, as well as the AMEs *aac(3)-IIa* and *aac(6’)-Ib-cr* in clade C2 ST131 carrying IncF plasmids, the majority of which came from patients in Singapore.

Previous work, mainly including isolates from North America, suggested that the *H*30-R/C1 clade of ST131 most commonly carry IncF type F1:A2:B20 plasmids and the *H*30-Rx/C2 clade are associated with IncF type F2:A1:B-plasmids.^17-19^ In this cohort, plasmid types were associated with different sub-lineages of ST131. For instance, IncF type F31:A4:B1 or F36:A4:B1 plasmids were most frequently seen in clade C2, whereas IncF type F2:A1:B-were only seen in a single clade C1 strain. These variations may reflect sampling from different geographical locations, rather than associations with specific *E. coli* lineages.

This study has some limitations. Enrolment into the clinical trial required susceptibility to piperacillin-tazobactam at the local testing laboratory, therefore bias may exist in the selection of strains. In addition, enrolment of patients into a clinical trial may preclude those with severe comorbidities or early mortality (prior to randomisation), it is possible that the *E. coli* were obtained from patients with less severe disease, which may be associated with less virulent strains.

The MERINO Trial is currently recruiting in an additional 6 countries (Italy, Turkey, Canada, South Africa, Lebanon and Saudi Arabia) and aims to report in 2018. It is anticipated that this current work can be augmented by these additional isolates and provide a global perspective on the molecular epidemiology of contemporary ESBL- or p-AmpC-producing *E. coli* causing bloodstream infections.

## Acknowledgements

We would like to acknowledge all the members of the study teams from the recruiting sites. **Royal Brisbane & Women’s Hospital**: Tiffany Harris-Brown, Penelope Lorenc, John McNamara. **Princess Alexandria Hospital**: Neil Underwood, Jared Eisenmann, James Stewart, Andrew Henderson. **National University Hospital**: Jaminah Ali, Donald Chiang. **Tan Tock Seng Hospital**: Soh Siew Hwa, Yvonne Kang, Ong Siew Pei, Ding Ying. **North Shore Hospital**: Umit Holland. **Monash Health:** Tony Korman, Infectious Disease Registrars 2015-2017

## Sequence data

Raw sequence reads and associated meta-data can have been uploaded to NCBI (Bioproject Accession number: PRJNA398288).

## Transparency declarations

PH and SAB have spoken at an educational event sponsored by Pfizer. BR has consulted for Mayne Pharma and Merck. PAT has received research support from GSK, Shionogi, Sanofi-Pasteur and Janssen in the last twelve months. DLP has received honoraria for advisory board participation and speaking at events sponsored by Achaogen, Merck and GlaxoSmithKline. All other authors declare no conflicts of interest.

## Contributions of authors

PH wrote the first and final drafts and undertook the laboratory work. AW and HMZ helped with the whole genome sequencing and NLBZ, LR and SB undertook the genomic data analysis. All other authors are site investigators for the trial and helped to recruit patients and collect bacterial isolates. DLP is the chief investigator for the MERINO study and conceived the concept for the paper with PH, NLBZ, LR and SB. All authors contributed to the writing of the paper and have approved the final version.

## Funding

This project was supported by funding from the Pathology Queensland Study, Education and Research Committee (SERC), the National University Hospital Singapore (NUHS) Clinician Researcher Grant, the Australian Society of Antimicrobials (ASA), the International Society for Chemotherapy (ISC) and the National Health and Medical Research Council (NHMRC) of Australia (GNT1067455). PH is supported by the Royal College of Pathologists of Australasia (RCPA) Foundation Postgraduate Research Fellowship and an Australian Postgraduate Award (APA) from the University of Queensland. SAB is supported by an NHMRC Career Development Fellowship (GNT1090456). MAS is supported by an NHMRC Senior Research Fellowship (GNT1106930).

